# A Genomics Resource for Genetics, Physiology, and Breeding of West African Sorghum

**DOI:** 10.1101/2020.06.03.132217

**Authors:** Jacques M. Faye, Fanna Maina, Eyanawa A. Akata, Bassirou Sine, Cyril Diatta, Aissata Mamadou, Sandeep Marla, Sophie Bouchet, Niaba Teme, Jean-Francois Rami, Daniel Fonceka, Ndiaga Cisse, Geoffrey P. Morris

**Affiliations:** Department of Agronomy, Kansas State University, Manhattan, KS, USA.; Institut Sénégalais de Recherches Agricoles, Centre d’Étude Régional pour l’Amélioration de l’Adaptation à la Sécheresse, Thiès, Sénégal.; Institut National de la Recherche Agronomique du Niger, Niamey Niger.; Institut Togolaise de Recherche Agronomique, Togo.; Institut d’Economie Rurale, Mali.; AGAP, Univ Montpellier, CIRAD, INRA, Montpellier SupAgro, Montpellier, France.; CIRAD, UMR AGAP, Thies, Senegal.

## Abstract

Local landrace and breeding germplasm is a useful source of genetic diversity for regional and global crop improvement initiatives. Sorghum (*Sorghum bicolor* L. Moench) in West Africa has diversified across a mosaic of cultures and end-uses, and along steep precipitation and photoperiod gradients. To facilitate germplasm utilization, a West African sorghum association panel (WASAP) of 756 accessions from national breeding programs of Niger, Mali, Senegal, and Togo was assembled and characterized. Genotyping-by-sequencing was used to generate 159,101 high-quality biallelic SNPs, with 43% in intergenic regions and 13% in genic regions. High genetic diversity was observed within the WASAP (π = 0.00045), only slightly less than in a global diversity panel (π = 0.00055). Linkage disequilibrium decayed to background level (*r*^*2*^ < 0.1) by ~50 kb in the WASAP. Genome-wide diversity was structured both by botanical type, and by populations within botanical type, with eight ancestral populations identified. Most populations were distributed across multiple countries, suggesting several potential common gene pools across the national programs. Genome-wide association studies of days to flowering and plant height revealed eight and three significant quantitative trait loci (QTL), respectively, with major height QTL at canonical height loci *Dw3* and *SbHT7.1*. Colocalization of two of eight major flowering time QTL with flowering genes previously described in US germplasm (*Ma6* and *SbCN8*) suggests that photoperiodic flowering in WA sorghum is conditioned by both known and novel genes. This genomic resource provides a foundation for genomics-enabled breeding of climate-resilient varieties in West Africa.

---

Crop production in many developing countries is limited by biotic and abiotic factors that reduce food supplies to smallholder farmers in semi-arid areas. With an increasing worldwide underfed population along with environmental changes, there is a need in more rapidly developing locally adapted varieties to increase crop productivity (Foley *et al.*, 2011; Tilman *et al.*, 2011; Mundia *et al.*, 2019). Genetic studies contribute to the development of adapted varieties to meet global food security and help provide enough genetic diversity suitable for efficient crop breeding (Jordan *et al.*, 2011). Diverse landrace germplasm harbors useful alleles for gene discovery and breeding, given their long history of adaptation to diverse environments (Meyer & Purugganan, 2013). However, African crop genetic diversity, particularly in West Africa (WA), are poorly characterized mainly due to lack of genomic resources and limited sampling of genetic resources available to the global scientific community.

Understanding genomic variation of local germplasm at a regional scale can help guide breeding. The availability of high-density markers evenly distributed throughout the genome is a prerequisite for understanding genetic diversity and genetic basis of adaptive traits. Recent advances in next-generation sequencing technologies and GBS have rendered possible the generation of high-density markers with affordable low cost (Elshire *et al.*, 2011; Poland *et al.*, 2012). These tools facilitate characterization of the genetic structure of local germplasm relative to global diversity. Historical recombination along with short to moderate linkage disequilibrium (LD) existing within a diversity panel greatly improve the mapping resolution to identify novel genes and novel natural variants at known genes in major crops (Huang *et al.*, 2010; Yano *et al.*, 2016; Cao *et al.*, 2016; Gapare *et al.*, 2017; Zhao *et al.*, 2019).

In sorghum, global reference diversity panels have been assembled (Grenier *et al.*, 2001; Deu *et al.*, 2006; Casa *et al.*, 2008; Upadhyaya *et al.*, 2009; Billot *et al.*, 2013; Brenton *et al.*, 2016) and used in genetic and breeding studies. However, at a regional scale these global reference panels have limited population sampling, thus limiting their potential use in regional association mapping studies. Regional diversity panels are useful genetic resources for capturing natural allelic variation existing in locally-adapted varieties (Leiser *et al.*, 2014; Sattler *et al.*, 2018). Favorable alleles for adaptation to various regional environmental conditions have been selected for over thousands of years, but they may not be easily accessible to breeders via global *ex situ* collections (Hammer & Teklu, 2008; Fu, 2017). The major West and Central African cereal crops such as sorghum and pearl millet need to be assembled into regional reference diversity panels that can be accessed for genetic, physiology, and breeding.

Genetic diversity of cultivated sorghum is high in West African (Doggett, 1988; Deu *et al.*, 1994; Folkertsma *et al.*, 2005). Despite the high genetic diversity in the West African germplasm, its accessibility to the regional and global scientific community is limited, particularly germplasm that was more recently collected or developed. Five botanical types in sorghum—bicolor, durra, guinea, caudatum and kafir and ten intermediate types—have been defined based on spikelet and grain morphology (Harlan & De Wet, 1972) and are associated with climate and geographic origin (Brown *et al.*, 2011). All botanical types are represented in West African landraces except for the kafir type. The guinea type is the most common and diverse in WA, possibly due to a second center of domestication (Folkertsma *et al.*, 2005). The durra type was first domesticated in Ethiopia before diffusing to arid regions of West Africa. The caudatum and durra-caudatum intermediate types are well represented in the west-central region, and used for grain yield improvement in breeding programs throughout West Africa (ISRA, 2005). However, little is known about the population structure of the germplasm across the countries, whether the germplasm of one country is distinct from other countries. Since country boundaries in West Africa cut across agro-ecological zones, some landrace germplasm may be more similar across countries than within a country.

Here, we report the assembly of the WASAP from the four West African countries (Mali, Niger, Senegal and Togo) and development of genome-wide SNP markers as genomic resources for genetics, physiology, and breeding. We determined the genome-wide SNP variation and population structure of the germplasm in relationship with previously genotyped West African *ex situ* collections and GDP. We also identified known maturity and plant height loci using GWAS on phenotype data collected under rainfed conditions. The study provides genomic resources and a better understanding of the population structure of the WA germplasm useful for genomics-assisted breeding.

## Materials and methods

### Plant materials

The WASAP was composed of 756 accessions assembled by breeders, physiologists, and geneticists in national agricultural research organizations (NARO) from four West African countries (Senegal, Mali, Togo, Niger) (Table 1; Supplemental Data S1). The panel includes working collections of the NARO sorghum improvement programs, predominantly landraces, but also locally-improved varieties and breeding lines. Based on *a priori* classification all four basic botanical types (bicolor, caudatum, durra, and guinea) are well represented in the WASAP, except the kafir type (n = 1). Many accessions were not classified morphologically into botanical classes (n = 230). Genetic diversity and structure of the WASAP were compared with the global diversity panel (GDP) that consists of 692 worldwide sorghum accessions (excluding accessions from Americas) with available sequencing data, including West African accessions in the GDP (hereafter named WAS-GDP) (Morris *et al.*, 2013; Lasky *et al.*, 2015). WASAP accessions were also compared to previously genotyped West African accessions from USDA-GRIN (hereafter named WAS-GRIN), originating from Niger (NiGRIN), Senegal (SnGRIN; including a few accessions from neighboring Gambia and Mauritania), and Nigeria (NGrGRIN) (Maina *et al.*, 2018; Olatoye *et al.*, 2018; Faye *et al.*, 2019).

**Table 1.**
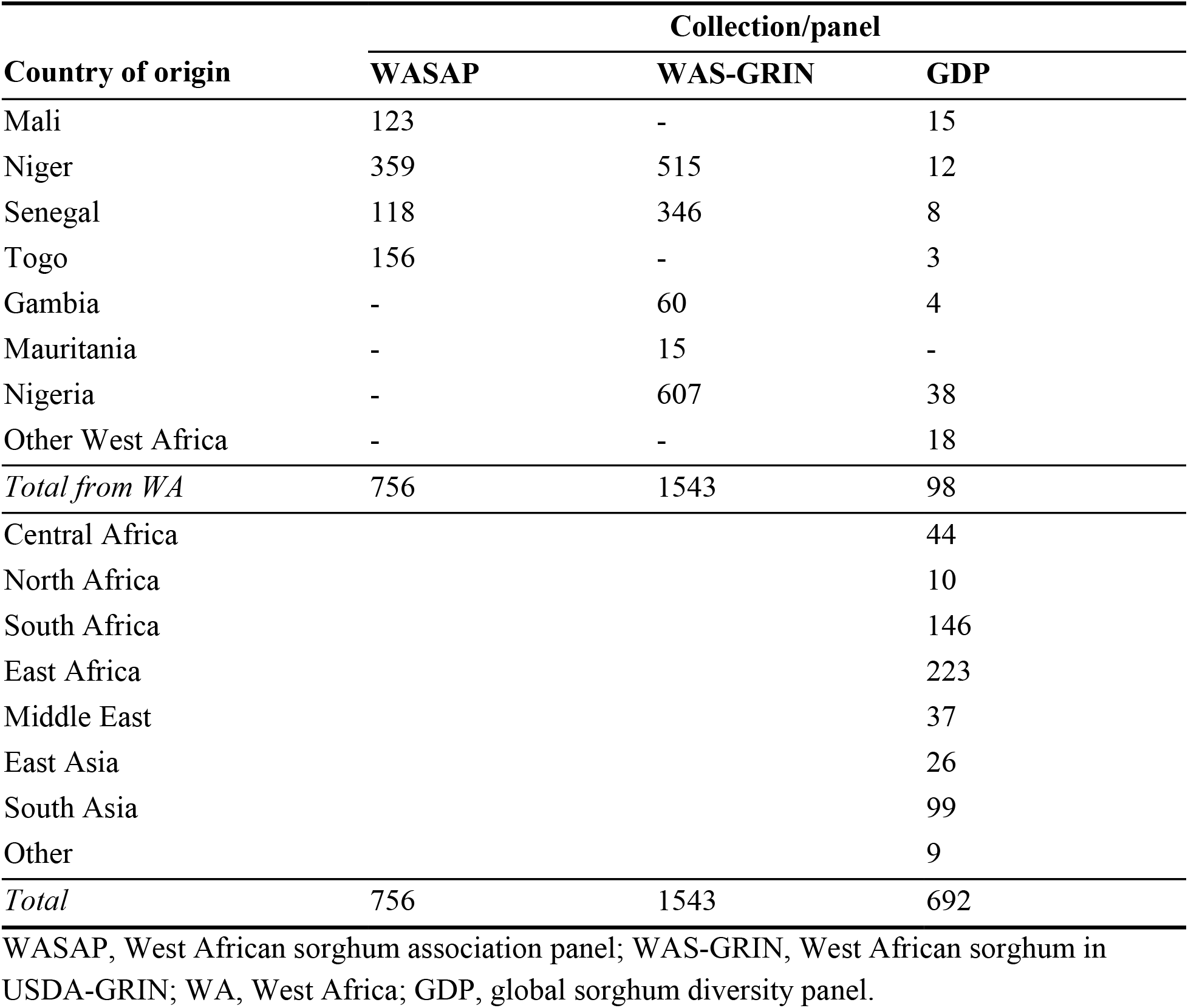
Number of accessions in each sorghum collection/panel used in this study.

| **Country of origin** | **Collection/panel** |  |  |
| --- | --- | --- | --- |
|  | **WASAP** | **WAS-GRIN** | **GDP** |
| Mali | 123 | - | 15 |
| Niger | 359 | 515 | 12 |
| Senegal | 118 | 346 | 8 |
| Togo | 156 | - | 3 |
| Gambia | - | 60 | 4 |
| Mauritania | - | 15 | - |
| Nigeria | - | 607 | 38 |
| Other West Africa | - | - | 18 |
| *Total from WA* | 756 | 1543 | 98 |
| Central Africa |  |  | 44 |
| North Africa |  |  | 10 |
| South Africa |  |  | 146 |
| East Africa |  |  | 223 |
| Middle East |  |  | 37 |
| East Asia |  |  | 26 |
| South Asia |  |  | 99 |
| Other |  |  | 9 |
| *Total* | 756 | 1543 | 692 |

WASAP, West African sorghum association panel; WAS-GRIN, West African sorghum in USDA-GRIN; WA, West Africa; GDP, global sorghum diversity panel.

### Genotyping-by-sequencing and SNP discovery

The WASAP was grown in the field at the Bambey research station in Senegal and leave tissue from five seedlings of each accession were pooled to extract genomic DNA using the MATAB (Mixed Alkyl trimethylammonium bromide) protocol. Genotyping-by-sequencing (GBS) was conducted following the method previously described (Elshire *et al.*, 2011). Briefly, GBS libraries were constructed in 96-plex and the restriction enzyme, *Ape*KI (New England Biolabs), was used to digest genomic DNA for complexity reduction. For quality control, a random well was left blank in each 96-well plate. Restriction cutting sites were ligated using barcoded adapters and ligated products were pooled together. Single-end sequencing was performed on Illumina HiSeq2500 at the University of Kansas Medical Center (Kansas, USA).

Illumina single-end sequence reads and GDP raw sequence data were processed together using the TASSEL GBS v2 pipeline (Glaubitz *et al.*, 2014). Sequence reads were trimmed to 64-bp and identical reads collapsed into tags. Tags were aligned to the sorghum reference genome v3.1 (Paterson *et al.*, 2009; McCormick *et al.*, 2018) using the Burrows-Wheeler Alignment (BWA) program (Li & Durbin, 2009). Single nucleotide polymorphism (SNP) markers were discovered using the *DiscoverySNPCallerPluginV2* of the TASSEL GBS v2 pipeline with *minimum locus coverage* (*mnLov*) of 0.1 and other parameters kept to default settings. A total of 546,133 SNPs were discovered. SNPs with more than 20% missing rate (n = 393,396 remaining SNPs), minor allele frequency (MAF) < 0.01 (n = 201,193 remaining SNPs), and monomorphic sites were filtering out. Biallelic SNPs were kept, resulting in a total set of 198,402 SNPs. This dataset was imputed using Beagle v4.1 (Browning & Browning, 2016) and filtered out again data with MAF < 0.01 to yield a final data set of 159,101 SNPs that was used for downstream analysis.

### Genome-wide SNP variation

Linkage disequilibrium (LD) was determined for the entire WASAP and for each country’s germplasm separately. SNP markers with MAF < 0.05 were filtered out before estimating LD based on the pairwise correlation coefficient (*r^2^*) among SNPs in a window of 500 kb using the PopLDdecay package (Zhang *et al.*, 2019). The *smooth.spline* function in the R program (R Core Team, 2016) was used to fit LD decay measured as the distance by which *r*^*2*^ decreased to half from its original value. The imputed dataset was functionally annotated to determine SNP effects on protein-coding genes using the snpEff program (Cingolani *et al.*, 2012). The structural location and functional class (synonymous, missense, or nonsense) of each SNP were determined based on the sorghum reference sequence. Genome-wide SNP distribution along chromosomes, minor allele frequencies, and pairwise nucleotide diversity were estimated using the VCFtools program (Danecek *et al.*, 2011). F_ST_ genetic differentiation in the WASAP was determined according to botanical type and country of origin using the pairwise F_ST_ method based on Weir and Cockerham weighted F_ST_ estimate in the VCFtools.

### Genetic structure analysis

The genome-wide SNP variation of the collection was assessed based on the principal component analysis (PCA) using the *snpgdsPCA* function of the R package SNPRelate (Zheng *et al.*, 2012). The accessions of WAS-GRIN and GDP were included in the analysis to robustly determine principal axes of genome-wide SNP variation and clustering of the WASAP along other germplasms. The combined dataset consisted of a subset of 103,871 (100K) high-quality SNP markers with a high level of polymorphism (MAF > 10%). The GDP accessions were used as a training set to calculate principal components and predict the genetic structure among the WASAP and WAS-GRIN accessions.

The number of ancestral populations and ancestry fractions in the WASAP were determined using ADMIXTURE (Alexander *et al.*, 2009). The original SNP dataset was LD-pruned to generate 60,749 SNPs using the function *--indep 50 10 2* in the PLINK 1.9 (Purcell *et al.*, 2007). ADMIXTURE was run with K = 2–20 and five-fold cross-validation was performed to identify the optimal K. We defined the optimal K as the minimum K where cross-validation error no longer decreased substantially. Accessions were assigned to subpopulations based on 0.7 membership threshold. The neighbor-joining distance-based clustering method was used to assess the genetic relationship among WASAP ancestral populations, WAS-GRIN, and GDP. The genetic distance matrix was generated based on the 100K SNPs using TASSEL 5 (Bradbury *et al.*, 2007) and the results were plotted using *ape* R package (Paradis *et al.*, 2004).

### Field phenotyping

Field phenotyping of a subset of 572 WASAP accessions (based on seed availability) was conducted at the Bambey Research Station, Centre National de Recherches Agronomiques (14.42°N, 16.28°W) in Senegal during the growing season of 2014. Two experiments, early-sown date at the beginning of the rainy season–normal planting date (Hiv1) and late-sown post-flowering drought-stressed–planted 30 days later (Hiv2), were performed. Each experiment had one replication. A randomized incomplete block design was performed with 30 blocks per experiment. Five check varieties were randomly assigned into each block of 19 genotypes. Days to flowering (DFLo) was measured as the day when 50% of plants within a plot flowered. Plant height (PH) was defined as the average distance from the soil to the top of three plants per plot.

### Statistical analysis of phenotypic data

Phenotypic variation was analyzed using the R program. The variance components were estimated by fitting the mixed linear model with random effects for all genotypes (G), environment (E), and G × E interaction effects using the *lme4* package (Bates et 2010). Broad sense heritability (H^2^) was estimated for each trait from the estimated genetic and residual variances derived from the mixed effect model, as follows:

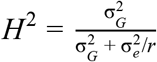

Where 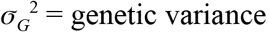, 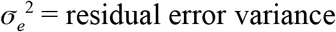, and *r* the number of environments. Phenotypic correlations among traits were calculated using the Pearson correlation of the *PerformanceAnalytics* package (Peterson *et al.*, 2014). BLUP values were calculated by combining data from environments. Tukey’s Honestly Significant Difference (TukeyHSD) in the *Agricolae* package (Mendiburu, 2009) was used to test the difference of genotype performance between environments.

### Genome-wide association studies

Genome-wide association studies (GWAS) were performed using the general-linear model (GLM) in the GAPIT R package (Lipka *et al.*, 2012) and multi-locus mixed-model (MLMM) in the *mlmm* package (Segura *et al.*, 2012). The MLMM stepwise regression model was used to account for background effects. A total of 130,709 SNPs with MAF > 0.02 were used for the GWAS analysis, as rare variants could contribute to phenotypic variation. The first three principal components and kinship matrix were generated from TASSEL 5 (Bradbury *et al.*, 2007) to account for polygenic background effects in the MLMM analysis. GWAS were performed for BLUP values of DFLo and PH across two environments. LD heatmaps of genomic regions surrounding GWAS QTL were constructed using the package *LD heatmap 0.99-4* package (Shin *et al.*, 2006). All figures were produced with the R program. The effect size and proportion of phenotypic variance explained by associated QTLs were determined using linear regression and ANOVA. Background effect was accounted for using ADMIXTURE ancestry fractions used as fixed effect covariates. Candidate gene colocalization with QTL was carried out using an *a priori* candidate genes/loci list, including known sorghum genes and orthologs of rice and maize for adaptive traits, previously described (Faye *et al.*, 2019). The Sorghum QTL Atlas (Mace *et al.*, 2019) was used to compare QTLs identified in the current study to QTLs from previous studies.

## Results

### Genome-wide SNP variation of the West African sorghum association panel

The GBS library sequencing yielded a total of ~258 million single-end sequencing barcoded reads. After trimming all reads down to 64 bp, ~4.5 million unique tags were obtained. A final data set of 159,101 high-quality SNPs was maintained after removing SNPs with >20% missing data, MAF < 0.01, and keeping only biallelic SNPs. The SNPs were distributed across the genome with a higher number of SNPs in pericentromeric regions relative to centromeric regions (Fig. 1A). We determined if the 159,101 GBS-SNPs have potential impacts on protein-coding sequences based on the sorghum reference sequence v.3.1. About 13% of the SNPs were found in genic regions, including 7,689 missense (72%), 411 (4%) nonsense, and 2,603 (24%) silent point mutations (Supplementary Fig. S1). Of these SNP, 43%, 22%, and 22% variants were located in intergenic, downstream, and upstream regions of genes, respectively.

**Fig. 1.**
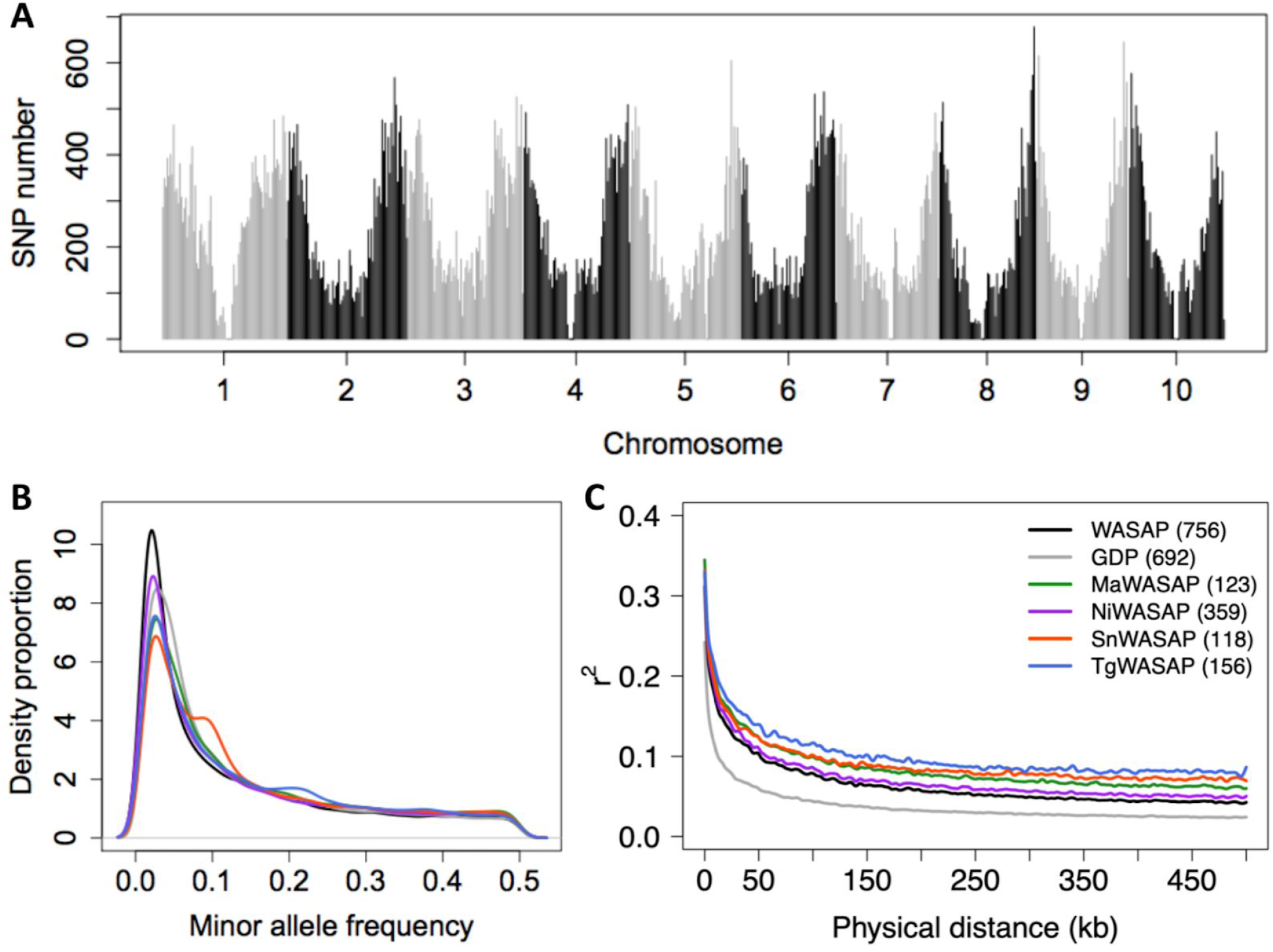
Genome-wide SNP variation in the WASAP and GDP. (A) Distribution of the SNP data across the 10 sorghum chromosomes in the WASAP. Minor allele frequency distribution and Linkage disequilibrium decay along the genome (C) of the SNP data in the whole WASAP, within country in WASAP–Mali (MaWASAP), Niger (NiWASAP), Senegal (SnWASAP) and Togo (TgWASAP), and in the global sorghum diversity panel (GDP).

Since four of five sorghum botanical types are represented in the WASAP, we hypothesized that it captures much of the genetic diversity found in the GDP. As predicted, the estimate of average pairwise nucleotide diversity (π) was only slightly less (18%) in the WASAP (0.00045) than in the GDP (0.00055). Little variation in π was observed among the four countries of origin (Niger: 0.00046, Mali: 0.00049, Senegal: 0.00050, Togo: 0.00047). The WASAP had a higher proportion of rare alleles (e.g., 0.01–0.05) than the GDP (Fig. 1B). Within the WASAP, the Senegal accessions had the lowest rare allele proportion and the highest intermediate allele proportion, followed by Togo and Mali. Linkage disequilibrium (LD) decayed to background level (*r*^*2*^ < 0.1) by ~50 kb in the WASAP versus ~15 kb in the GDP. LD decayed to background by ~60 kb, ~90 kb, ~90 kb, and ~160 kb in Niger, Senegal, Mali, and Togo accessions, respectively (Fig. 1C).

### Genetic differentiation by botanical types and geographic origin

Based on the hypothesis that sorghum genetic diversity is structured primarily by botanical type, we predicted high F_ST_ genetic differentiation would be observed among botanical types than among countries of origin in the WASAP. High F_ST_ genetic differentiation of 0.16 was observed among the six classes composing the majority of the panel, including guinea (G), caudatum (C), durra (D), bicolor (B), guinea-margaritiferum (Gm) types, and the intermediate form durra-caudatum (DC) (Supplemental Table S1). Surprisingly, high F_ST_ value was observed between DC and C (F_ST_ = 0.22) or DC and D (F_ST_ = 0.20). The F_ST_ value among the four countries of origin was moderate (F_ST_ = 0.09) (Supplemental Table S1).

Next we used PCA to characterize the genomic variation of the WASAP with respect to botanical type and origin, in comparison with WAS-GRIN and GDP accessions. The first two PCs explained a high proportion of genome-wide SNP variation (a combined 17%) and differentiated the caudatum, durra, guinea, and kafir accessions (Fig. 2A). We predicted that the majority of the WASAP accessions will be clustered with guinea, caudatum, and durra accessions from the same geographic origin in the GDP. The majority of the WASAP accessions overlapped with their corresponding types, guinea, durra, and caudatum clusters of GDP along the PC2. A substantial variation was explained by both PC3 vs. PC4 (7.3%) (Fig. 2B). The durra-caudatum intermediate types in the WASAP and WAS-GRIN clustered between durra, caudatum, and guinea clusters in the GDP.

**Fig. 2.**
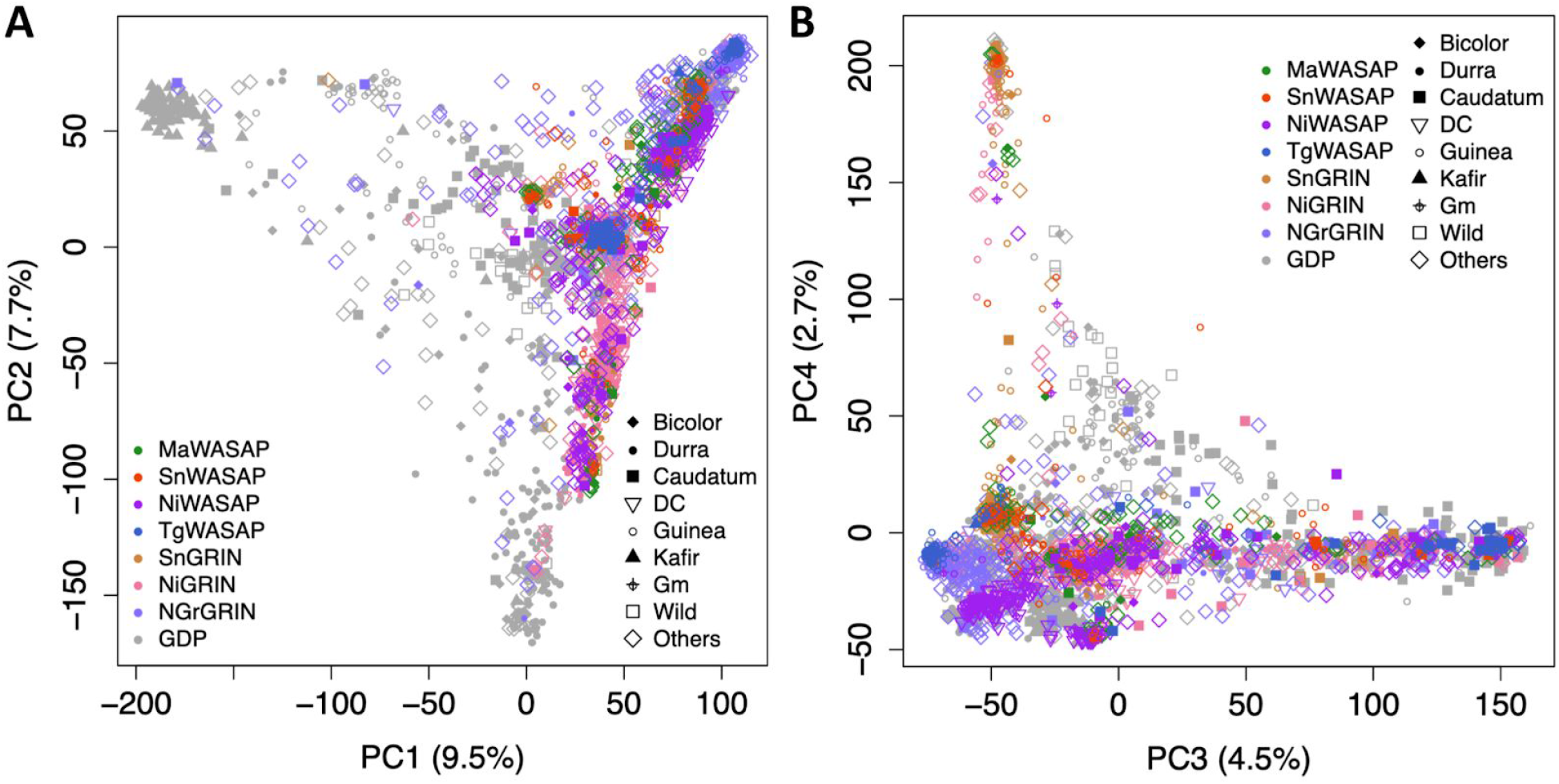
Principal component analysis of genome-wide SNP variation. Scatterplots of the first and second axes (A) and the third and fourth axes (B) of genome-wide SNP variation in the WASAP in relationship with other West African sorghums in GRIN and global sorghum diversity panel. The color-codes indicate country of origin for WASAP accessions (MaWASAP, Mali; NiWASAP, Niger; SnWASAP, Senegal and TgWASAP, Togo), the West African accessions in GRIN (SnGRIN, Senegal, Gambia and Mauritania; NiGRIN, Niger; NGrGRIN, Nigeria), and the global sorghum diversity panel (GDP). The symbols indicate botanical types where DC and Gm correspond to durra-caudatum intermediate and guinea-margaritiferum types, respectively.

### Ancestral fractions and population structure

To determine the ancestral populations and ancestry fractions for each accession, we used the Bayesian model-based method ADMIXTURE. Based on five-fold cross-validation error (CVE), the optimum number of ancestral populations was eight (Fig. 3A). The accessions classified morphologically as guinea (orange lower rug-plot) corresponded to three genetic groups (G-II, G-IV, G-VII) (Fig. 3B, Supplemental Data S1). The accessions classified morphologically as durra-caudatum intermediates (mostly from Niger; green lower rug-plot) corresponded to two genetic groups (G-V, G-VIII). Accessions classified morphologically as caudatum (blue lower rug-plot) corresponded to G-I. Using 0.7 ancestry fraction as a threshold, 71% of accessions could be assigned to a subpopulation, while 29% would be considered admixed. The greatest putative contribution to genetic admixture was from G-V (purple bars), with ancestry fraction present in all other subpopulations. F_ST_ among ancestral populations averaged 0.39, with a range of 0.25–0.61 (Supplemental Table S2).

**Fig. 3.**
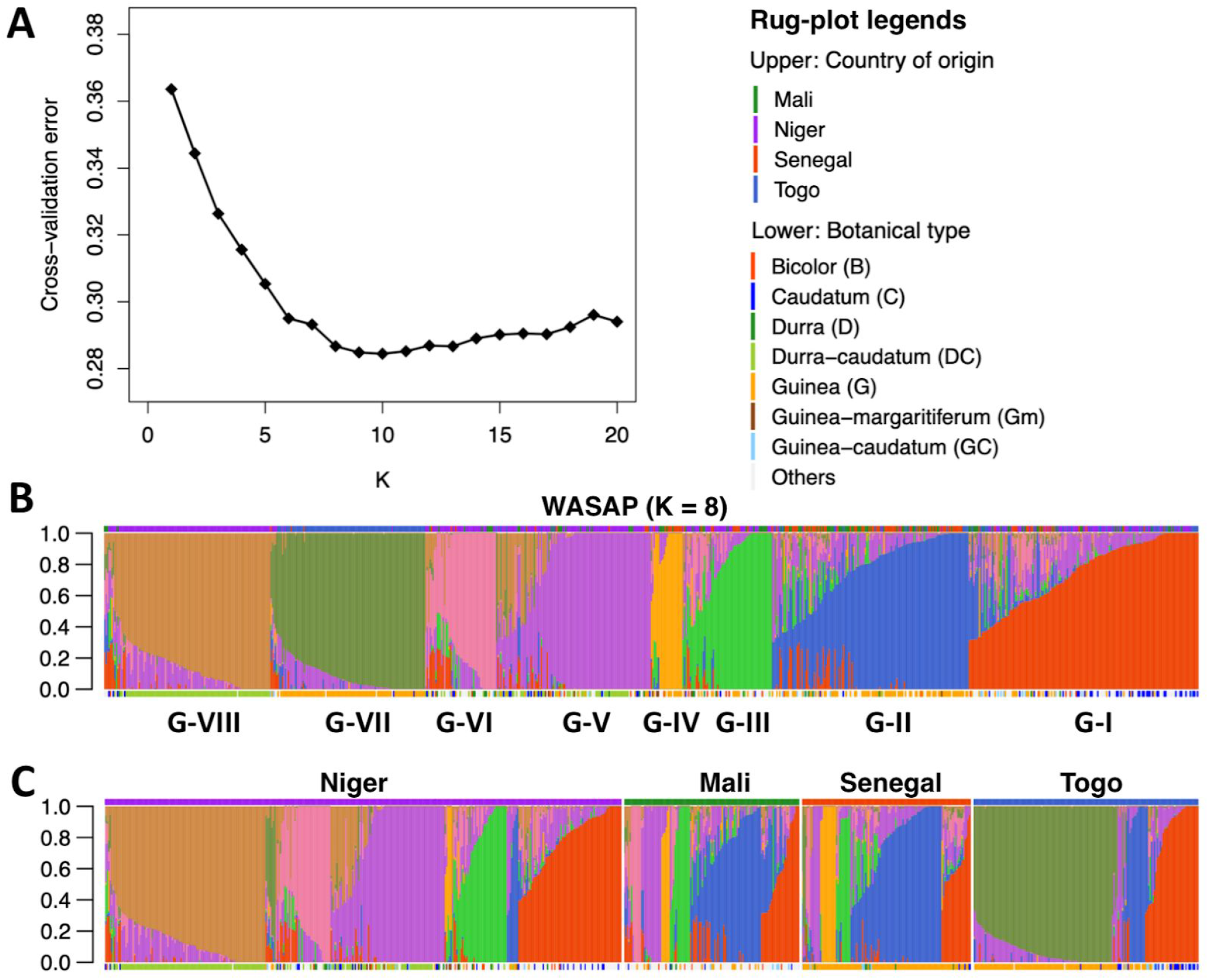
Genetic ancestry analysis of the WASAP. (A) Five-fold cross-validation error from the ADMIXTURE model using 60,749 SNPs for K = 2–20. Ancestral genetic groups of the WASAP at K = 8 ancestral populations (B) ordered by ancestry fraction and (C) ordered by country then by ancestry fraction. Vertical bars represent ancestry fraction from the eight ancestral populations (G-I to G-VIII) indicated with a different arbitrary color for each population (Note, ancestral population color-coding in the vertical bars does not correspond to the rug-plot color-coding). Upper rug-plots indicate countries of origin. Lower rug-plots indicate botanical type (“Others” include rare intermediate types and accessions of unknown botanical type). Ancestry fractions for each accession are in Supplemental Data S1.

Next, we considered the extent to which germplasm of each country is distinct from the germplasm of other countries (Fig. 3C). Each of the countries’ germplasm included multiple genetic groups. Most of the ancestral populations were found in each country, except in Togo where only three genetic subpopulations (G-I, G-II, G-VII) were clearly defined. The G-IV was specific to Senegal and Mali, while G-VI was specific to Niger and Mali. G-VII and G-VIII groups were specific to Togo and Niger, respectively. Neighbor-joining (NJ) analysis recapitulated the country-level ancestry structure (see color-coded tips) (Fig. 4 and Supplementary Fig. S2). Genetic similarities were observed between WASAP ADMIXTURE genetic groups and other West African sorghums (WAS-GRIN and WAS-GDP) from the same geographic origin. and GDP accessions according to botanical type and geographic origin. The West African sorghums clustered with their corresponding types in the GDP. The guinea and durra-caudatum accessions of West Africa clustered generally distinctly from the majority of GDP accessions.

**Fig. 4.**
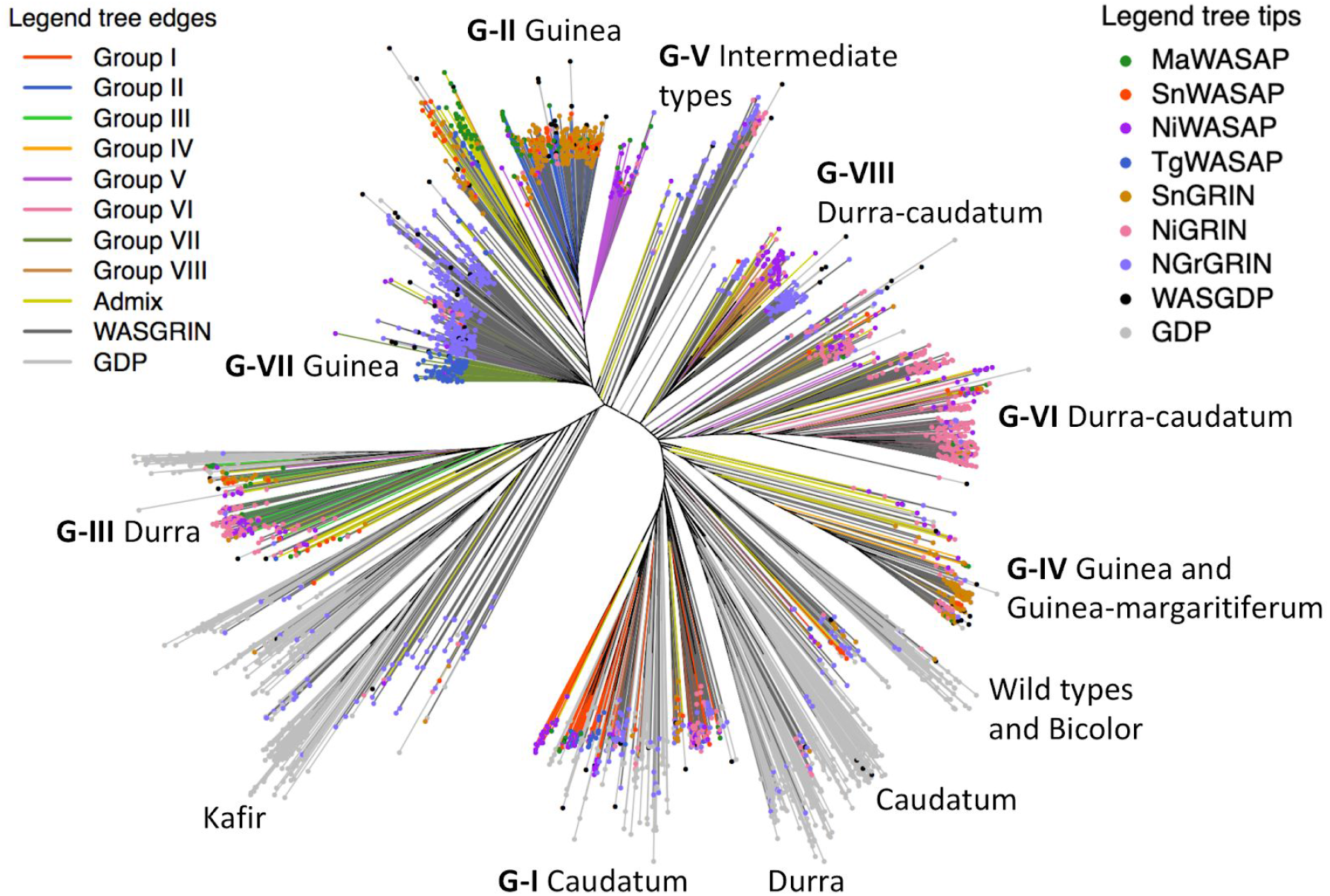
Neighbor-joining analysis of the WASAP. Clustering of the WASAP accessions (MaWASAP, Mali; NiWASAP, Niger; SnWASAP, Senegal and TgWASAP, Togo) in relationship with other West African sorghums in GRIN (SnGRIN, Senegal, Gambia and Mauritania; NiGRIN, Niger; NGrGRIN, Nigeria) and global sorghum diversity panel (GDP). The color-coding of the tree edges is based on the ADMIXTURE ancestral populations (G-I to G-VIII, including admixed accessions) of the WASAP. The edges in yellow, dark gray, and light gray represent admixed WASAP accessions (< 0.6 ancestry fraction), WASGRIN accessions, and GDP accessions, respectively. The color-coding of the tree tips indicate accessions origin, with black tips indicating West African sorghum accessions in the GDP (WASGDP).

### Phenotypic variation in the WASAP

A total of 572 accessions of the WASAP and five check lines were evaluated for agronomic traits under rainfed conditions. We hypothesized that the phenotypic variation in the WASAP is supported by genetic effect, appropriate for mapping with GWAS. Large phenotypic variation was observed for both DFLo (21%) and PH (35%). Large genotypic (G) variation and genotype by environment (G × E) interaction effects were observed (Table 2). High broad-sense heritability (*H*^*2*^) was high across the two environments, with values ranging from 0.86 for PH to 0.93 for DFLo (Table 2). DFLo were significantly higher in Hiv1 (TukeyHSD *p* < 0.0001; Supplemental Fig. S3), but significantly correlated, based on the 1:1 ratio, between Hiv1 and Hiv2 (r^2^ = 0.75, *p* < 0.0001) (Supplemental Fig. S4A). The relationship holds within country of origin, for instance for accessions from Togo (r^2^ = 0.77, *p* < 0.0001) and Senegal accessions (r^2^ = 0.78, *p* < 0.0001) (Supplemental Fig. S4B). DFLo and PH were significantly correlated in each environment (Supplemental Fig. S5).

**Table 2.** Descriptive statistics and phenotypic variation across early (Hiv1) and late (Hiv2) planting date experiments under rainfed conditions.

| Trait | Range | Mean ±<br>SD | CV (%) | G (%) | E (%) | GxE (%) | $H^2$ |
| --- | --- | --- | --- | --- | --- | --- | --- |
| DFLo | 40–128 | 78 ± 15 | 21 | 79 | 1 | 9 | 0.93 |
| PH (cm) | 65–410 | 214 ± 75 | 35 | 57 | 3 | 20 | 0.86 |

DFLo, days to flowering; PH, plant height; SD, Standard deviation; CV, coefficient of variation; G, genotype and E, environment variances; $H^2$ , broad-sense heritability.

### Genome-wide association studies for flowering time and plant height

We assessed the effectiveness of the genome-wide SNP data for genetic dissection of complex quantitative traits using GWAS. For DFLo BLUPs, the GLM naive model identified many significant associations at Bonferroni correction 0.05 (Fig. 5A). A leading QTL was identified near *Ma6/ Ghd7* candidate gene between SNPs S6_651847 (top association in the region, ~45 kb away) and S6_699843 (within gene). A second leading QTL was identified between S9_54917833 and S9_54968379 at 43 kb and 4 kb from *SbCN8* gene, respectively (Supplemental Table S3). The GLM showed inflation of *p*-values with many significantly associated SNPs. The MLMM model identified eight QTL at Bonferroni threshold (3.8 × 10^−7^). Some of these QTLs colocalized with known candidate genes *Ma6* at 45 kb (S6_651847) and *SbCN8* at 381 kb (S9_55345348) (Table 3). The QTL near *Ma6*, S6_651847, was the top peak in the region in both GLM and MLMM and was at one gene away from *Ma6* (Fig. 5C and D). LD between the QTL, S6_651847 and SNPs near/within *Ma6* locus was moderate (Fig. 5E). After controlling for the population structure, the association of S6_651847 had an estimated effect size of 29 days and PVE of 25% (Table 3; Fig. 5F).

**Table 3.** Quantitative-trait loci associated with days to flowering BLUPs across early and late planting date experiments using the MLMM.

| QTL SNP<sup>a</sup> | MLMM<br>*p*-value | MAF | Effect<br>size | PVE<sup>b</sup><br>(%) | MLM<br>*p*-value | Distance to<br>locus (kb) | Locus<br>name |
| --- | --- | --- | --- | --- | --- | --- | --- |
| S4_57407080 | <10<sup>-12</sup> | 0.02 | 10 | 7 | 0.002 |  |  |
| S5_18513685 | <10<sup>-12</sup> | 0.10 | 12 | 9 | 0.0008 |  |  |
| S8_2206437 | <10<sup>-10</sup> | 0.04 | 28 | 11 | 0.0001 |  |  |
| S9_55345348 | <10<sup>-9</sup> | 0.04 | 15 | 9 | 0.0005 | 381 | *SbCN8* |
| S6_651847 | <10<sup>-8</sup> | 0.25 | 29 | 25 | <10<sup>-11</sup> | 45 | *Ma6* |
| S5_42404204 | <10<sup>-8</sup> | 0.12 | 6 | 16 | 0.0001 |  |  |
| S10_51083132 | <10<sup>-7</sup> | 0.28 | 7 | 5 | 0.03 |  |  |
| S2_11509143 | <10<sup>-7</sup> | 0.04 | 5 | 0.3 | 0.6 |  |  |

<sup>a</sup> Digit before and after underscore indicates chromosome number and SNP position on the genome, respectively; <sup>b</sup> ADMIXTURE ancestry fractions at K = 8 were used as fixed effect covariates; BLUP, best linear unbiased prediction; MLMM, multi-locus mixed-linear model; QTL, quantitative-trait loci; MAF, minor allele frequency; GLM, general linear model; PVE, proportion of variance explained; MLM, mixed-linear model.

**Fig. 5.**
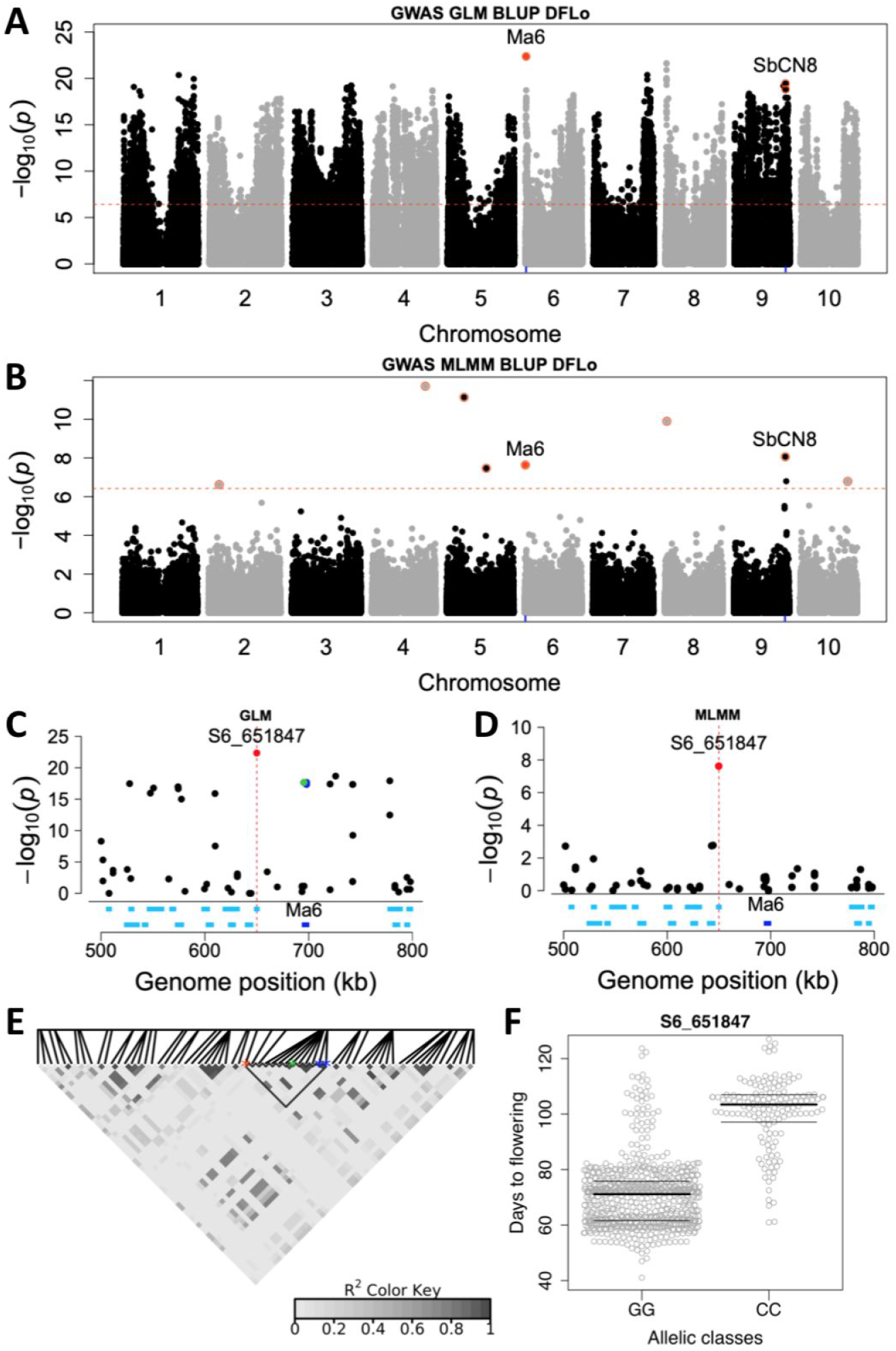
GWAS for days to flowering (DFLo) under rainfed conditions. Manhattan plots of DFLo based on (A) the GLM and (B) the MLMM. The horizontal red dashed line represents the Bonferroni significance threshold at 0.05. The rug plots indicate the position of colocalizing candidate genes, *Ma6* and *SbCN8* with QTLs. Regional Manhattan plot of a 150 kb region on chromosome 6 around the QTL S6_651847 that colocalizes with *Ma6* from (C) GLM and (D) MLMM. The green and blue peaks are SNP QTLs at 160 bp from and within *Ma6*, respectively. The dark blue segment indicates the genomic position of *Ma6*. (E) LD heatmap of a 150 kb region surrounding the QTL S6_651847. The red, green, and blue asterisks indicate the S6_651847, SNPs at 160 bp from *Ma6* and within *Ma6*, respectively. (F) Days to flowering across planting dates by allelic classes of the QTL S6_651847.

For PH, the GLM naive model identified several associations (Fig. 6A). The top association, S7_56232413 overlapped with the height QTL *qHT7.1* (Li *et al.*, 2015; Bouchet *et al.*, 2017). A second QTL was identified between SNPs S7_59955806 (top association in the region) and S7_59896357 located at 66 kb from the *Dw3 a priori* candidate gene. The MLMM identified three QTLs at the Bonferroni threshold (Fig. 6B; Supplemental Table S4). A putative SNP QTL, S7_59400476 was identified 421 kb away from *Dw3*, though below the Bonferroni threshold. After accounting for confounding population structure, the MLMM QTLs still had significant estimated effect sizes and contributed to high PVE for PH (Supplemental Table S4). LD between S7_59955806 and S7_5940047 was high, but weak between these SNP QTLs and a variant in *Dw3* locus (Fig. 6E). Alleles at both SNP QTLs were associated with height differences of accessions across planting dates (Fig. 6F and G). The association of S7_5940047 with PH, which had the highest allelic effect estimate (73 cm) and PVE (41%), was confirmed using MLM (*p* < 10^−13^) (Supplemental Table S4).

**Fig. 6.**
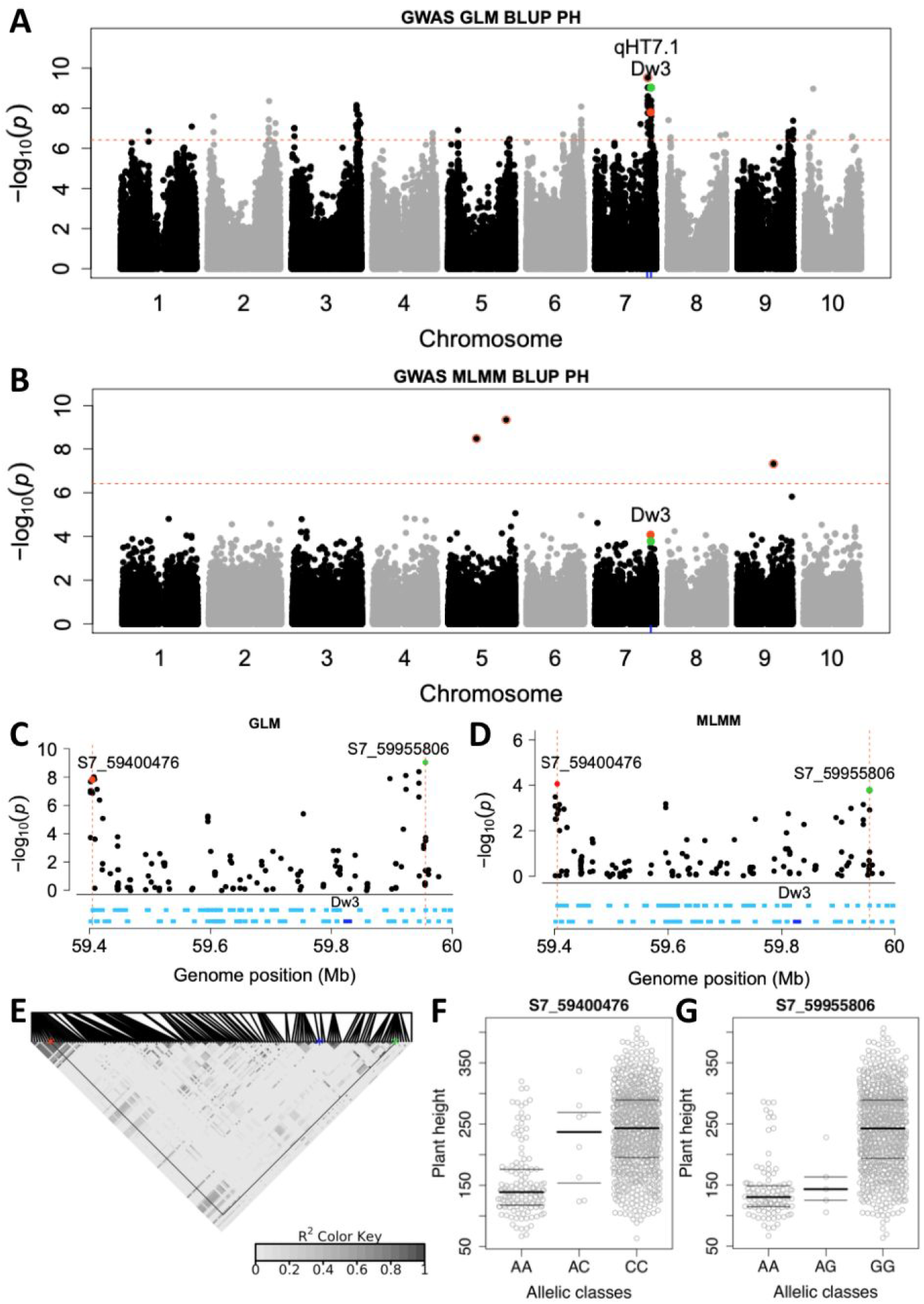
GWAS for plant height (PH) under rainfed conditions. Manhattan plots of PH based on (A) the GLM and (B) the MLMM. The horizontal dashed line represents the Bonferroni significance threshold at 0.05. Rug plots on chromosome 7 indicate the position of the candidate gene, *Dw3* and *qPH7.1*. Regional Manhattan plot of a 600 kb region on chromosome 7 surrounding the QTL between S7_59400476 and S7_59955806 that colocalizes with *Dw3* from GLM and (D) MLMM. The red and green peaks are top SNPs in MLMM and GLM, respectively. The dark blue segment indicates the genomic position of *Dw3*. (E) LD heatmap of genomic region between SNPs S7_59400476 and S7_59955806. The red, green, and blue asterisks indicate the S7_59400476, S7_59955806, and a SNP within *Dw3*, respectively. Days to flowering across planting dates by allelic classes of SNP QTLs (F) S7_59400476 and (G) S7_59955806.

## Discussion

### Developing a high-quality regional genomic resource

In the present study, we assembled 756 sorghum accessions from West African sorghum germplasm and generated high-density genome-wide SNP dataset. We demonstrated that this genome-wide SNP dataset is of sufficient quality for genomic and quantitative genetic analyses suitable for crop improvement through genomics-assisted breeding. The strict data filtering criteria used before and after genotype imputation provided a final dataset with reduced number of SNPs (n = 159,101) many of which have impacts on protein-coding sequences (Supplementary Fig. S1A).

The quality control of the SNP dataset matched our expectations as the F_ST_, PCA, and neighbor-joining analyses (Figs. 2, 4 and Supplemental Fig. S2) confirmed the expected structure of sorghum by botanical type and geographic region (Morris *et al.*, 2013; Lasky *et al.*, 2015; Bouchet *et al.*, 2017; Wang *et al.*, 2019). The validity of the SNP dataset was further confirmed based on GWAS with the identification of QTLs (Figs. 5 and 6) that colocalized with known candidate loci, *Ma6* and *SbCN8* (Yang *et al.*, 2014; Murphy *et al.*, 2014) for days to flowering and *qHT7.1* and *Dw3* (Multani *et al.*, 2003; Li *et al.*, 2011, 2015) for plant height. Although regional diversity panels could limit confounding effects due to population structure in GWAS, in this case the strong population structure in the WASAP appeared to increase false positive associations in the GLM (Brachi *et al.*, 2011). However, using the stepwise regression in MLMM though helped to control the inflation of *p*-values observed in the GLM (Fig. 5 and 6).

Resolution of GWAS depends on linkage disequilibrium (LD) decay across the genome (Slatkin, 2008; Korte & Farlow, 2013). LD decay range was generally short in the WASAP and in the GDP (Fig. 1C) compared to the long LD range of the GDP reported in previous studies. In sorghum, studies have reported variation in LD decays from short (10–15 kb) to moderate (50–100 kb) (Hamblin *et al.*, 2005; Bouchet *et al.*, 2012) to about 150 kb in some studies with higher density of markers and larger population size (Morris *et al.*, 2013; Mace *et al.*, 2013b). The shorter LD range observed in the GDP in this study could be explained by the exclusion of North American breeding lines that share many common haplotypes (Klein *et al.*, 2008). Within the WASAP, the longer LD range in Togo accessions and shorter LD in Niger accessions (Fig. 1C) are consistent with the limited number of genetic subpopulations observed in Togo compared to the strong genetic structure in Niger (Fig. 3C).

### Insights into hierarchical population structure in West African sorghum

Sorghum genetic studies have identified population structure by botanical type and geographic region at regional scale (Deu *et al.*, 2006; Bouchet *et al.*, 2012; Morris *et al.*, 2013; Wang *et al.*, 2019) and at country level in the Senegal and Niger germplasm (Deu *et al.*, 2008; Faye *et al.*, 2019). The genetic diversity of the WASAP was structured by botanical type within each country of origin (Fig. 3 and 4). This finding is congruent with the F_ST_ analysis (Supplemental Table S1) where botanical type and country of origin contributed to high and moderate genetic differentiation, respectively.

The guinea type in the WASAP was split into three major subgroups (Fig. 4). One subgroup was formed by Senegal and Mali accessions, a second subgroup formed by Togo accessions (which clustered with Nigeria accessions), and a third subgroup that was more related to durra and durra-caudatum types, formed predominantly by Senegal accessions. This third group (G-IV) clustered with guinea margaritiferum accessions from Niger and was more related to bicolor and wild sorghums in the GDP. Previous studies (Deu *et al.*, 2006) have found four guinea subgroups; however, these subgroups included guinea sorghums from southern Africa, which were not represented in the WASAP. There were not significant genetic differences between WASAP and WAS-GRIN populations from the same geographic origin, suggesting that the globally-accessible GRIN collections are representative of current germplasm in West African. In contrast, overall genetic differences were observed between West African accessions and GDP where few subpopulations were formed almost entirely by West African sorghum accessions (e.g., G-VI and G-VII, Fig. 4), highlighting the distinctiveness of West African germplasm.

The high genetic diversity observed within each of the four countries of the WASAP (Fig. 3C) is relevant for breeding programs in the region. Within the WASAP, all eight ancestral populations were found in the germplasm of each country, except in the Togo germplasm, which appeared to be less diverse. Altogether, the genetic diversity of WA sorghum germplasm is hierarchically structured by botanical type and subpopulation within botanical type, with many subpopulations distributed across countries.

### Suitability of genomic resources for GWAS

To establish the utility of our genome-wide SNP dataset for GWAS, we carried out GWAS for flowering time and plant height and demonstrated colocalization of QTLs with known genes from previous studies. While flowering time genes and natural variants have been characterized in the US sorghum (Murphy *et al.*, 2014; Casto *et al.*, 2019), the genetic basis of the substantial photoperiodic flowering time variation in West African sorghum is not yet known (Bhosale *et al.*, 2012). The QTL S6_651847 near *Ma6* (*Gdh7*) (Murphy *et al.*, 2014) highly contributed to the proportion of phenotypic variation of flowering time. This QTL was mapped in both GLM and MLMM (Fig. 5C and D), indicating that *Ma6* might be a major flowering time gene in the WA sorghum germplasm. Only two SNPs, which were in low LD with the QTL S6_651847 (Fig. 5E), were found within *Ma6* locus. Several of the other identified QTLs overlapped with flowering time QTLs found in other studies based on the sorghum QTL Atlas (Mace *et al.*, 2019) (Supplemental Table S5).

Our findings are consistent with the hypothesis that a substantial oligogenic component underlies flowering time variation in WA sorghums, suggesting that photoperiodic flowering could be selected using markers from large effect QTLs. The two MLMM height QTLs (S5_30001948, and S9_38942669) accounted for 13.3% and 20.9%, respectively of height variation (Supplemental Table S4) but were not colocalized (within 500 kb) with known height genes. The major effect QTLs from this study would need to be validated with multi-year phenotypic data and other approaches, such as multi-parent linkage mapping and nearly-isogenic lines. Following validation, these height and flowering QTLs could be useful for marker-assisted selection to recover locally-adaptive plant types, across botanical types within West Africa or during prebreeding with exotic trait donors (Yohannes *et al.*, 2015). Given the positive findings for height and flowering time, we anticipate this resource will be suitable for characterization and mapping of other complex traits of interest to West African breeding programs.

### Implications for sorghum improvement in West Africa and globally

This study demonstrates that breeding populations in each of the four countries in West Africa harbor sufficient genetic diversity. The hierarchical population structure observed in the WASAP at country level suggests the existence of multiple ancestral populations within the country but similar across West Africa (Folkertsma *et al.*, 2005; Deu *et al.*, 2008; Sagnard *et al.*, 2011). The increased kinship within each subpopulation enables the implementation of genomic selection within individual subpopulation and/or a subpopulation across breeding programs. Otherwise, the strong population structure could lead to biased prediction accuracy as a result from allele frequency differences among subpopulations (Isidro *et al.*, 2015).

This SNP dataset could be used by the genetics community in genomic prediction, genetic diversity and haplotype analyses, and GWAS to serve global sorghum breeding, especially in combination with other regional genomic resources. Historically, West African sorghum germplasm has been useful for global sorghum breeding, including durra and caudatum accessions that were sources of yellow endosperm and drought tolerance for US breeding programs (Rosenow & Dahlberg, 2000). Whole genome-resequencing would complement the GBS-SNP dataset to more easily identify putative causal variants for adaptive traits in the West African germplasm (Bellis *et al.*, 2020). The development of trait-predictive markers could facilitate more rapid breeding of locally- and regionally-adapted sorghum varieties for the diverse stakeholders of West Africa.

## Abbreviations

BLUP: best linear unbiased prediction
CVE: cross-validation error
DFLo: days to flowering
GBS: genotyping-by-sequencing
GDP: global sorghum diversity panel
GLM: general linear model
GWAS: genome-wide association studies
LD: linkage disequilibrium
MLMM: multi-locus mixed linear model
PH: plant height
PVE: proportion of phenotypic variance explained
PW: panicle weight
QTL: quantitative trait loci
SNP: single nucleotide polymorphism
USDA-GRIN: United States Department of Agriculture, Germplasm Resources Information Network
WA: West Africa
WASAP: West African sorghum association panel.

## Data Availability Statement

The SNP and phenotypic datasets generated from this study are available in the Dryad Data Repository under accession [*to be added after acceptance*].

## Conflict of interest

The authors declare no conflict of interest.

## Author Contributions

Conceived and managed the study: GPM, DF, NC, AM, JFR. Contributed the genetic materials: NT, AM, EAA, CD, NC. Collected the data: SM, SB, FM, JMF, BS, EAA. Analyzed and visualized the data: JMF. Wrote the manuscript: JMF, GPM. All authors edited and approved the manuscript.

## Acknowledgments

This study is made possible by the support of the American People provided to the Feed the Future Innovation Lab for Collaborative Research on Sorghum and Millet through the United States Agency for International Development (USAID) under Cooperative Agreement No. AID-OAA-A-13-00047. The contents are the sole responsibility of the authors and do not necessarily reflect the views of USAID or the United States Government. The study was conducted using resources at the Integrated Genomics Facility and Beocat High-Performance Computing Cluster at Kansas State University. This study is contribution 20-318-J of the Kansas Agricultural Experiment Station.

## Supplemental Figures

**Supplemental Fig. S1.**
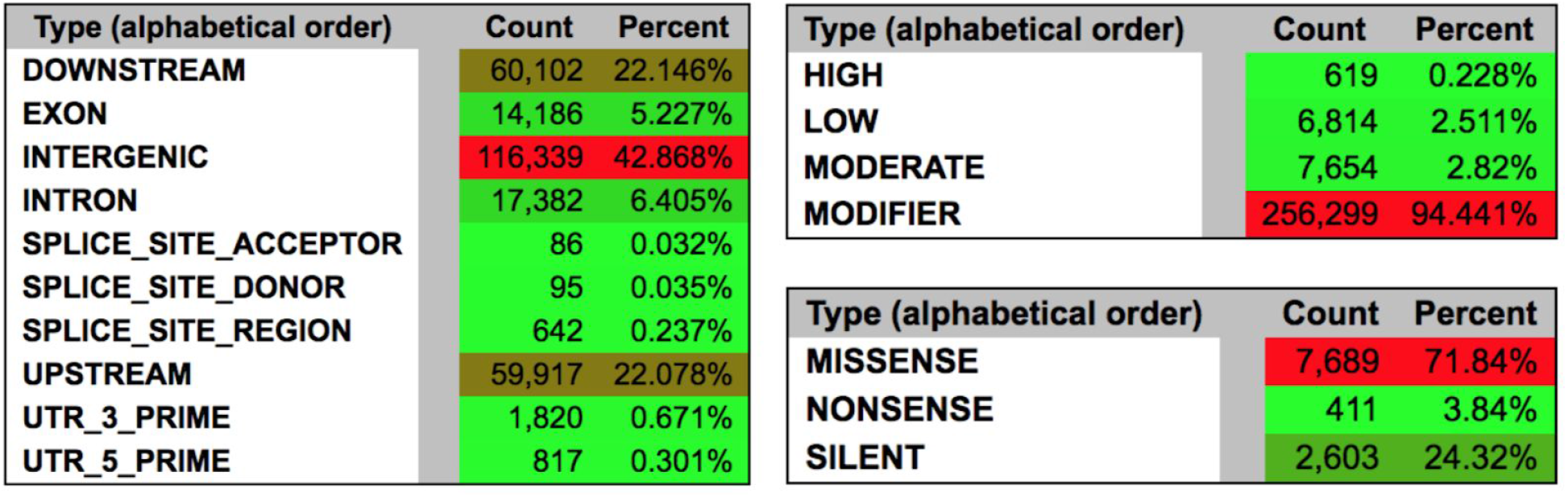
Genome-wide single nucleotide polymorphisms in the WASAP. (A) SNP annotation effect of 159,101 SNPs with MAF > 0.01.

**Supplemental Fig. S2.**
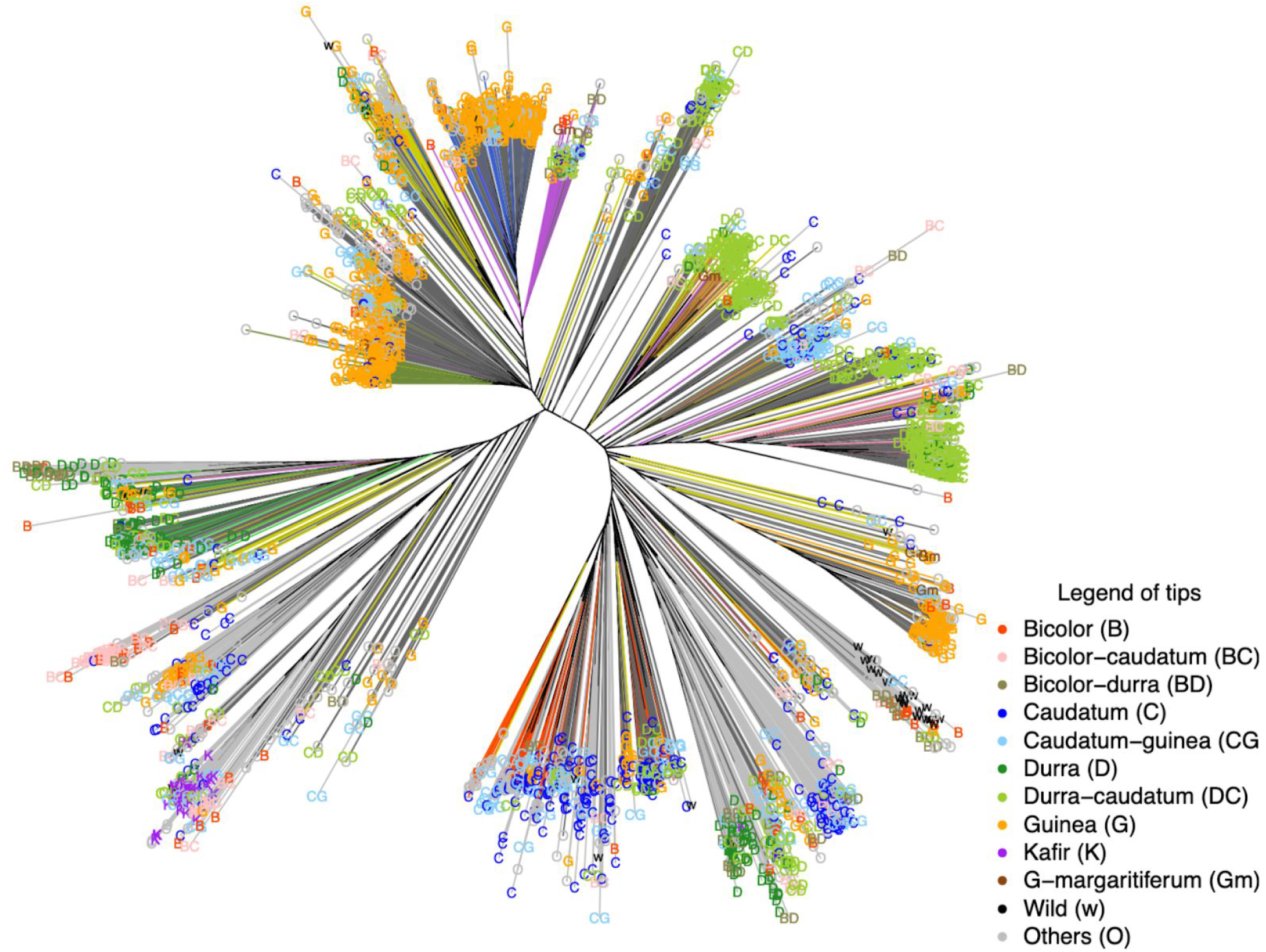
Unrooted Neighbor-joining tree of WASAP ancestral populations in relationship with other West African sorghums in GRIN (WAS-GRIN) and the global sorghum diversity panel (GDP). The tree edges and tips are color-coded based on the ADMIXTURE ancestral populations and the accessions origin, respectively. Tree edges in yellow, darkgray, and gray represent admixed (< 0.6 ancestry fraction) accessions in the WASAP, WAS-GRIN (SnGRIN, Senegal, Gambia and Mauritania; NiGRIN, Niger; NGrGRIN, Nigeria), and global sorghum diversity panel (GDP), respectively.

**Supplemental Fig. S3.**
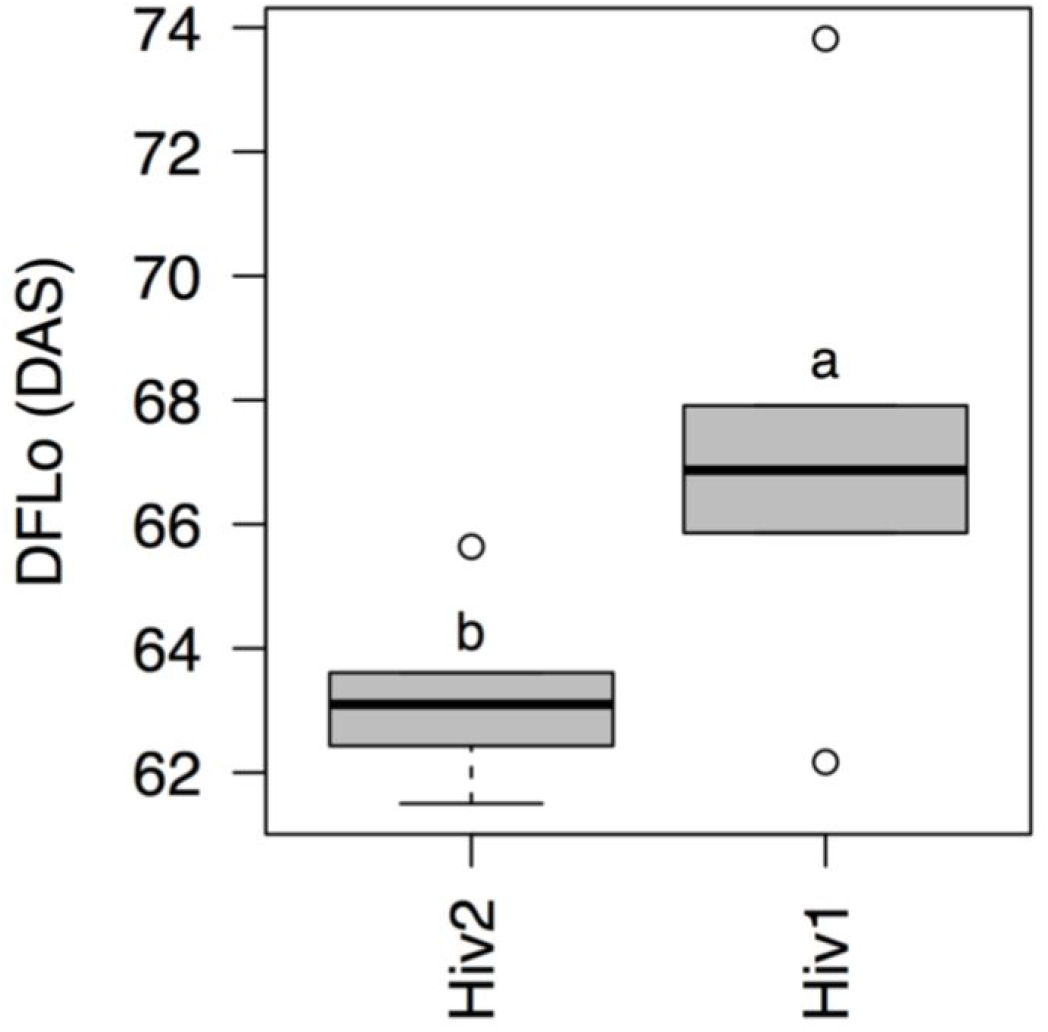
Average performance values for days to flowering (DFLo) of check varieties within each experiment under rainfed conditions. Different letters (e.g., a and b) indicate a significant difference between early (Hiv1) and late (Hiv2) planting date experiments based on the Tukey’s Honest Significant Difference test, *p* value < 0.0001. DAS means the number of days after sowing (planting).

**Supplemental Fig. S4.**
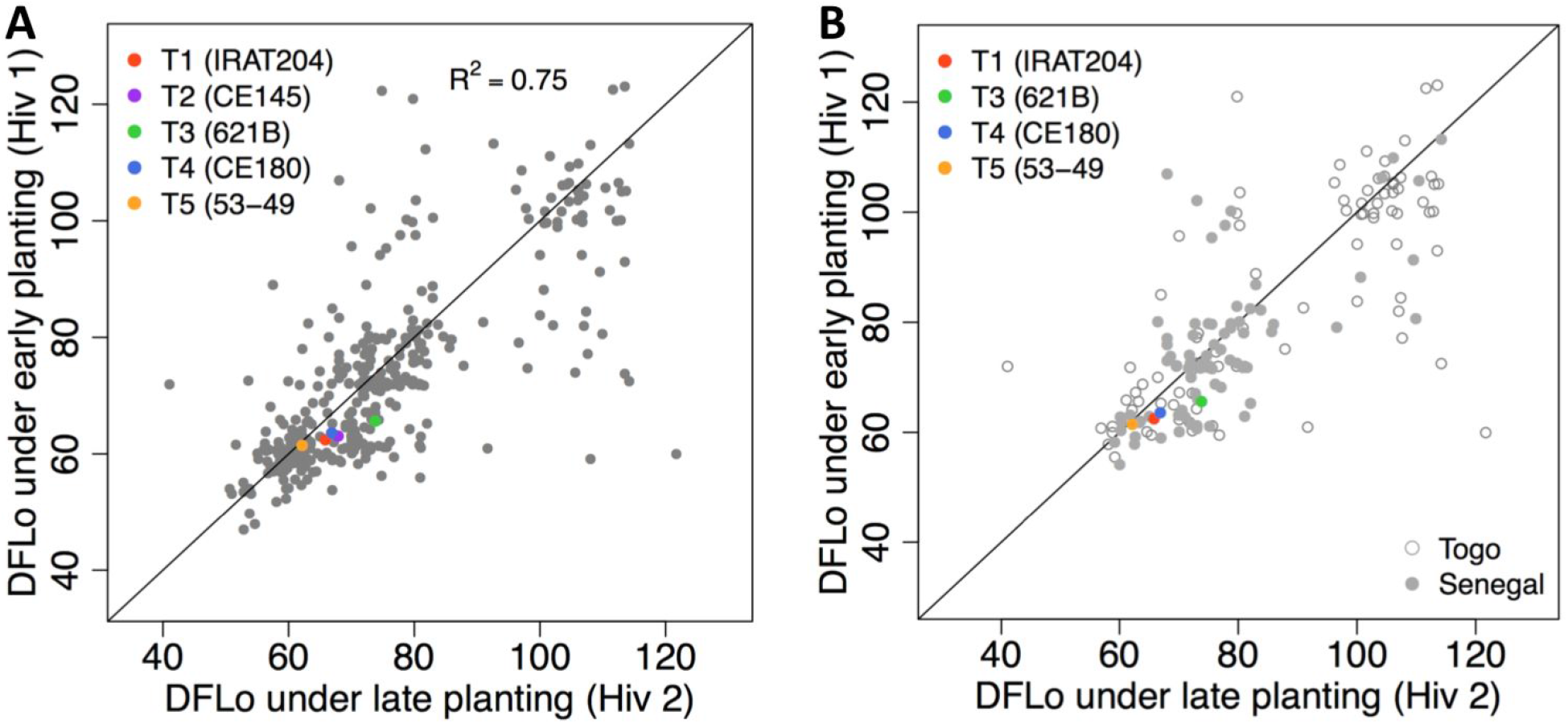
Flowering time differences of accessions between early (Hiv1) and late (Hiv2) planting date experiments under rainfed conditions. (A) Correlation for days to flowering (DFLo) between Hiv1 and Hiv2 within the whole WASAP. (B) Correlations for DFLo between Hiv1 and Hiv2 within Togo accessions (open circles) and within Senegal accessions (closed circles). The color-coded dots indicate the check varieties.

**Supplemental Fig. S5.**
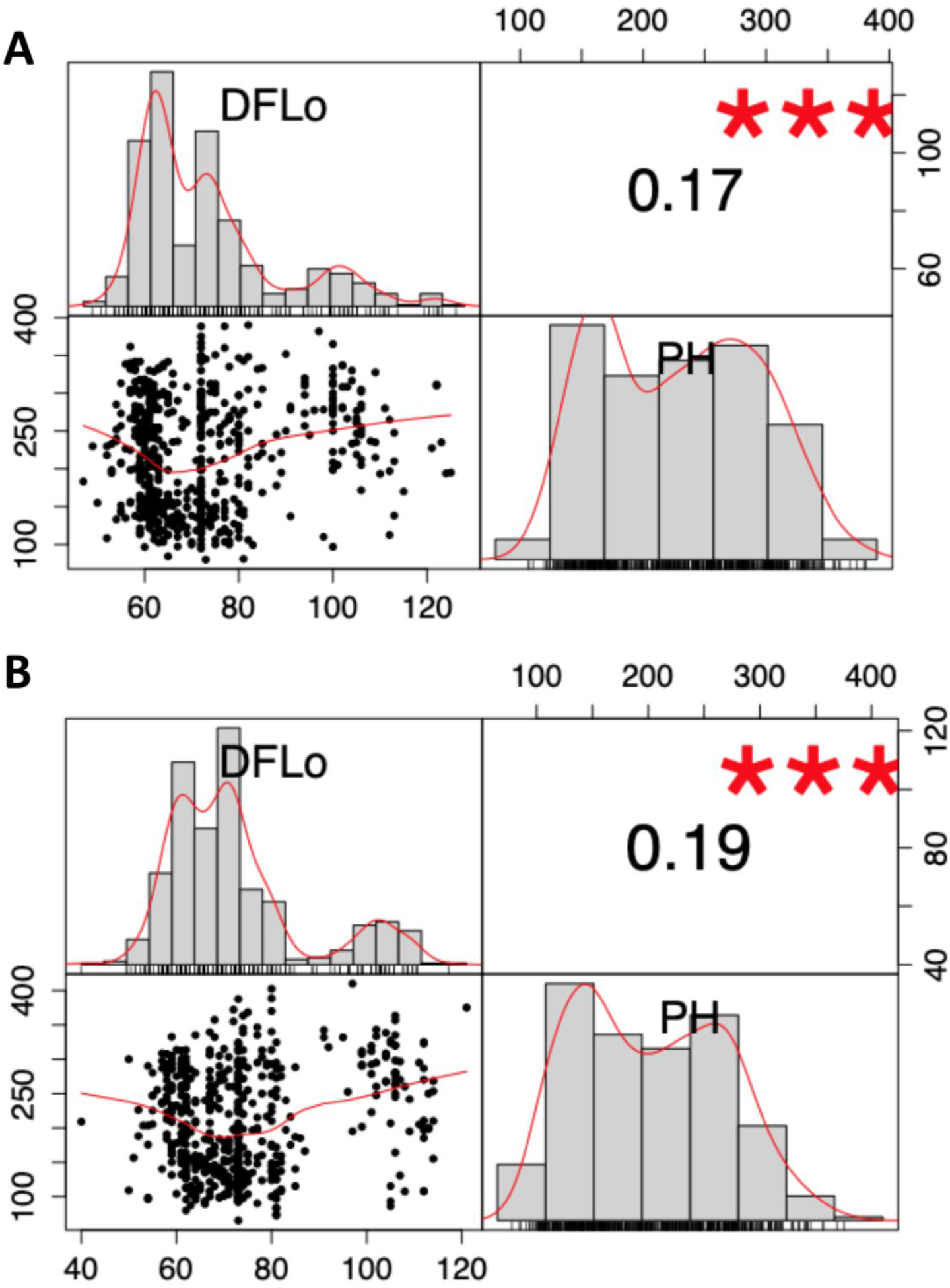
Phenotypic correlations for DFLo, days to flowering and PH, plant height within (A) early (Hiv1) and (B) late (Hiv2) planting date experiments.

## Supplemental Tables

**Supplemental Table S1.** Pairwise weighted F_ST_ genetic differentiation among botanical types and countries of origin.

| **(A) $F_{ST}$ among botanical types** |  |  |
| --- | --- | --- |
| Botanical type | | $F_{ST}$ |
| Bicolor | Durra | 0.02 |
| Bicolor | Caudatum | 0.05 |
| Bicolor | Guinea | 0.11 |
| Bicolor | Gm | 0.07 |
| Bicolor | DC | 0.18 |
| Durra | Caudatum | 0.11 |
| Durra | Guinea | 0.14 |
| Durra | Gm | 0.11 |
| Durra | DC | 0.20 |
| Caudatum | Guinea | 0.15 |
| Caudatum | Gm | 0.14 |
| Caudatum | DC | 0.22 |
| Guinea | Gm | 0.04 |
| Guinea | DC | 0.18 |
| Gm | DC | 0.17 |
| *Average* |  | 0.16 |
| **(B) $F_{ST}$ among countries of origin** |  |  |
| Country of origin | | $F_{ST}$ |
| Mali | Niger | 0.06 |
| Mali | Senegal | 0.01 |
| Mali | Togo | 0.12 |
| Niger | Senegal | 0.08 |
| Niger | Togo | 0.14 |
| Senegal | Togo | 0.12 |
| *Average* |  | 0.09 |

DC, durra-caudatum types; Gm, guinea margaritifera.

**Supplemental Table S2.** Pairwise weighted F_ST_ among ADMIXTURE ancestral populations.

| Populations | G-I | G-II | G-III | G-IV | G-V | G-VI | G-VII | Total $F_{ST}$ |
| --- | --- | --- | --- | --- | --- | --- | --- | --- |
| G-II | 0.36 |  |  |  |  |  |  |  |
| G-III | 0.35 | 0.43 |  |  |  |  |  |  |
| G-IV | 0.49 | 0.50 | 0.54 |  |  |  |  |  |
| G-V | 0.31 | 0.25 | 0.43 | 0.61 |  |  |  |  |
| G-VI | 0.34 | 0.44 | 0.32 | 0.54 | 0.43 |  |  |  |
| G-VII | 0.43 | 0.34 | 0.51 | 0.57 | 0.34 | 0.51 |  |  |
| G-VIII | 0.39 | 0.36 | 0.44 | 0.54 | 0.26 | 0.42 | 0.41 | 0.39 |

Accessions with >0.6 ancestry fractions for the given genetic group were included in $F_{ST}$ analysis.

**Supplemental Table S3.** Quantitative trait loci near *Ma6* and *SbCN8* candidate genes associated with days to flowering BLUPs using the GLM.

| QTL<sup>a</sup> | *P*-value | MAF | Position (kb) | Locus name |
| --- | --- | --- | --- | --- |
| S6_651847 | < 10<sup>-23</sup> | 0.25 | 45 | *Ma6* |
| S6_697299 | < 10<sup>-18</sup> | 0.24 | 160 bp | *Ma6* |
| S6_699842 | < 10<sup>-18</sup> | 0.22 | within | *Ma6* |
| S6_699843 | < 10<sup>-18</sup> | 0.23 | within | *Ma6* |
| S9_54917833 | < 10<sup>-20</sup> | 0.25 | 43 | *SbCN8* |
| S9_54968379 | < 10<sup>-19</sup> | 0.26 | 4 | *SbCN8* |

<sup>a</sup> Digit before and after underscore indicates chromosome number and SNP position on the genome, respectively; BLUP, best linear unbiased prediction; GLM, general linear model; QTL, quantitative trait locus; MAF, minor allele frequency.

**Supplemental Table S4.**
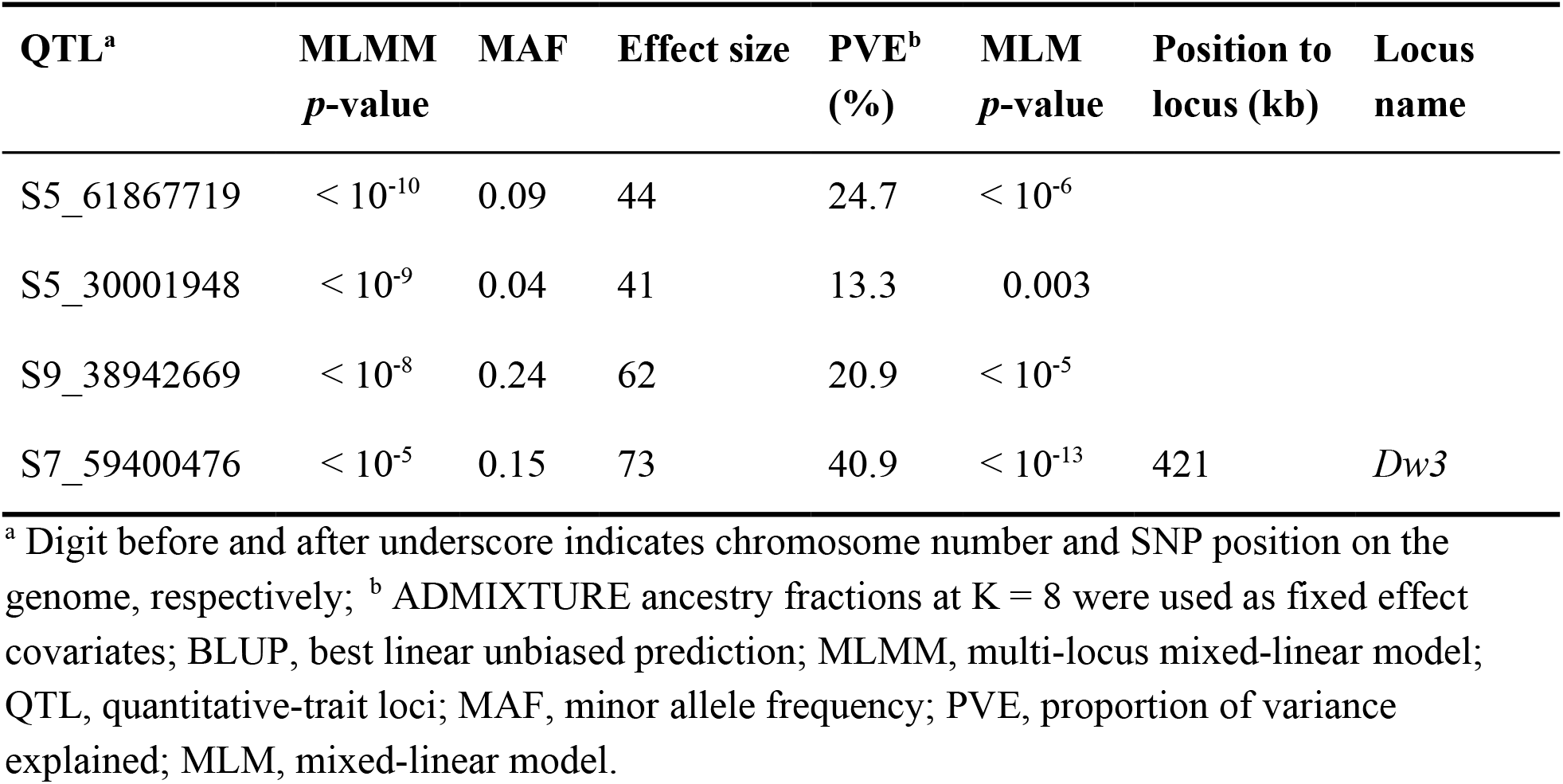
Quantitative-trait loci associated with plant height BLUP across early and late planting date experiments using the MLMM

| QTL<sup>a</sup> | MLMM<br>*p*-value | MAF | Effect size | PVE<sup>b</sup><br>(%) | MLM<br>*p*-value | Position to<br>locus (kb) | Locus<br>name |
| --- | --- | --- | --- | --- | --- | --- | --- |
| S5_61867719 | < 10<sup>-10</sup> | 0.09 | 44 | 24.7 | < 10<sup>-6</sup> |  |  |
| S5_30001948 | < 10<sup>-9</sup> | 0.04 | 41 | 13.3 | 0.003 |  |  |
| S9_38942669 | < 10<sup>-8</sup> | 0.24 | 62 | 20.9 | < 10<sup>-5</sup> |  |  |
| S7_59400476 | < 10<sup>-5</sup> | 0.15 | 73 | 40.9 | < 10<sup>-13</sup> | 421 | *Dw3* |

<sup>a</sup> Digit before and after underscore indicates chromosome number and SNP position on the genome, respectively; <sup>b</sup> ADMIXTURE ancestry fractions at K = 8 were used as fixed effect covariates; BLUP, best linear unbiased prediction; MLMM, multi-locus mixed-linear model; QTL, quantitative-trait loci; MAF, minor allele frequency; PVE, proportion of variance explained; MLM, mixed-linear model.

**Supplemental Table S5.** List of GWAS QTLs using MLMM, excluding those at *Ma6*, *SbCN8*, and *Dw3* overlapping with published QTLs from other studies based on the sorghum QTL Atlas.

| GWAS QTL<sup>a</sup> | QTL ID | LG:start–end | Original Reference |
| --- | --- | --- | --- |
| Days to flowering QTLs |  |  |  |
| S4_57407080 | *QDTFL4.21* | 4:56.1–61.6 | (Felderhoff *et al.*, 2012) |
| S8_2206437 | *QDTFL8.5* | 8:1.9–2.7 | (Wang *et al.*, 2014) |
|  | *QDTFL8.8* | 8:2.3–2.7 | (Mace *et al.*, 2013a) |
| S9_55345348 | *QDTFL9.16* | 9:50.4–57.9 | (Lin *et al.*, 1995) |
|  | *QDTFL9.15* | 9:50.4–57.9 | (Feltus *et al.*, 2006) |
|  | *QDTFL9.33* | 9:56.1–59.1 | (Zhang *et al.*, 2015) |
| S10_9523248 | *QDTFL10.10* | 10:7.6–9.7 | (Wang *et al.*, 2014) |
| S10_51083132 | *QDTFL10.26* | 10:42.1–51.9 | (Sangma, 2013) |
| Plant height QTLs |  |  |  |
| S5_61867719 | *QHGHT5.11* | 5:62.3–62.9 | (Bouchet *et al.*, 2017) |

<sup>a</sup> Digit before and after underscore indicates chromosome number and SNP position on the genome, respectively; QTL, quantitative trait locus; MLMM, multi-locus mixed-linear model; LG, linkage group; SNP, single nucleotide polymorphism

## Notes

### Competing Interest Statement

The authors have declared no competing interest.

